# Complex temperature profiles recorded through antisense co-transcriptional sensitivity

**DOI:** 10.64898/2026.11.10.765395

**Authors:** Jin Xu, Govind Menon, Chaocheng Guo, Miguel Montez, Xianhao Jin, Minglei Yang, Maria Sindalovskaya, Svenja Reeck, Susan Duncan, Eduardo Mateo-Bonmatí, Congyao Xu, Geng-Jen Jang, Yueying Zhang, Jixian Zhai, Yiliang Ding, Caroline Dean

## Abstract

Plants integrate widely fluctuating temperature over seasons to regulate their development. In vernalization, *FLOWERING LOCUS C* (*FLC*)^1,2^ is repressed by cold-induced antisense (*COOLAIR*) transcription and epigenetically silenced by Polycomb (PRC2) epigenetic switching^3^. Here, we show how *COOLAIR* co-transcriptional dynamics are central to integrating different facets of the natural temperature profile over different timescales. Constant cold alters *COOLAIR* RNA polymerase II processivity, splicing and polyadenylation site choice with distinct kinetics over days to weeks. In contrast, fluctuations below freezing at the same mean temperature caused extensive premature termination and chromatin retention of *COOLAIR*. High-resolution chromatin contact mapping showed *COOLAIR* premature termination altered local chromatin topology. This enhanced *FLC* promoter-gene body contacts, coincident with H3K4me1 loss and reduced *FLC* transcription. Multiple features of *COOLAIR* co-transcriptional sensitivity therefore act as thermosensory antennae, integrating distinct facets of winter temperature through local chromatin regulation, to robustly repress *FLC* in natural conditions.

## INTRODUCTION

Organisms use environmental cues to coordinate developmental transitions with favourable seasonal conditions. For temperature cues, the wide fluctuations over hourly and daily timescales, are often greater than the average changes over a season, so how gene regulatory systems integrate different aspects of temperature variation remains poorly understood. Vernalization, the acceleration of flowering by prolonged winter cold, provides a tractable system for investigating this problem.

In *Arabidopsis*, vernalization involves progressive repression of *FLOWERING LOCUS C* (*FLC*), a major floral repressor^1,2^. During cold, *FLC* transcription declines while Polycomb Repressive Complex 2 (PRC2) establishes epigenetic silencing through deposition of histone H3 lysine 27 trimethylation (H3K27me3)^4^. This silencing is maintained after plants return to warm conditions^5^. In parallel to Polycomb-mediated regulation, antisense transcription across *FLC* produces noncoding RNAs collectively termed *COOLAIR* that contribute to *FLC* repression during cold^3,6–9^. The antisense-associated repression pathway can occur over fast and slow timescales, whereas the PRC2-dependent silencing at *FLC* is a low probability switching mechanism, and thus always slow^3,10^.

The mechanism by which *COOLAIR* represses *FLC* has been investigated through mutations and natural variants. These affect *COOLAIR* splicing and polyadenylation, altering *FLC* expression and flowering^11–13^. Previous studies in warm-grown plants have also linked *COOLAIR* regulation to the local chromatin state at *FLC*. *COOLAIR* co-transcriptional processing and histone demethylation feedback on each other to affect productive *FLC* transcription and the establishment of Polycomb silencing^11–16^. These local chromatin changes include enhanced contact between the *FLC* promoter and its 3’ region^3,17^, and formation of *COOLAIR* RNA condensates over the *FLC* locus, termed ‘clouds’^18^. RNA structure of specific *COOLAIR* isoforms also affects chromatin association and *FLC* expression^8^.

A serendipitous finding during Arabidopsis vernalization experiments in natural and climatically distinct field sites, was a strong induction of *COOLAIR* expression the first night temperatures dipped below zero^19^. This was confirmed in chamber experiments comparing fluctuating and constant cold conditions that had the same mean temperature. *COOLAIR* hyper-expression correlated with stronger *FLC* repression^19^, but not with increased H3K27me3 accumulation^3^. We have therefore continued to investigate how sustained cold and brief fluctuations below freezing affect *COOLAIR* and *FLC* transcription. We developed targeted FLEP-seq (tFLEP-seq), which uses long-read sequencing of chromatin-associated RNA to resolve splicing and transcript 3’ ends, and combined it with Pol II profiling, RNA imaging, genetic perturbations and chromatin-contact mapping. Our findings show how the sensitivity of *COOLAIR* transcriptional processing enables plants to integrate different facets of the natural temperature profile over the prolonged timescales of winter. Coupling noncoding co-transcriptional sensitivity to environmental variation may be a general mechanism for robust gene regulation.

## Results

### Cold induces temporally distinct *COOLAIR* co-transcriptional changes across the locus

Gradual reduction in *FLC* expression during cold exposure results from reduced productive transcriptional initiation^20^. Counterintuitively, this is associated with increased co-transcriptional *FLC* splicing, which is influenced by antisense (*COOLAIR*) transcription^20^. We therefore investigated the transcriptional dynamics on the *COOLAIR* strand using native elongating transcript sequencing^21^ (NET-seq) over the vernalization time course analysed previously for *FLC* transcription and splicing (Figure S1A, B). RNA polymerase II (Pol II) levels on the *COOLAIR* strand were higher in the cold (Figure S1A, C), and its spatial distribution revealed unexpected features. On the *FLC* strand, Pol II accumulated predominantly at the *FLC* 3’ end^20^, as observed for many *Arabidopsis* genes^22,23^. On the *COOLAIR* strand, however, Pol II accumulated at two sites, indicative of slowed elongation or stalling (Figure S1A). The first, at the 3’ end of the proximal *COOLAIR* isoforms, increased and broadened after one week of cold (Figure S1A). The second, which we term the *COOLAIR* stalling site (CSS), occurs two-thirds into the *COOLAIR* transcription unit within the large intron, at a site not previously linked to polyadenylation (Figure S1A). The CSS lies ∼350 bp from the H3K27me3 nucleation region and close to the vernalization response element^24^. This is conserved in Brassicaceae *FLC*^9^ and deletion of the CSS region affects *FLC* regulation^24,25^. Cold reduced the proportion of Pol II signal beyond the CSS, in the region corresponding to the *FLC* promoter and *COOLAIR* class II polyadenylation sites (Figure S1D), indicative of cold-enhanced termination at the CSS. Thus, cold induced qualitative changes in Pol II occupancy on the *COOLAIR* strand that differed markedly from those on the *FLC* strand.

We reasoned that these temperature-sensitive transcriptional dynamics would be linked to changes in co-transcriptional processing. However, the short-read (Illumina) NET-seq data could not resolve differential splicing or polyadenylation. We therefore developed targeted FLEP-seq (tFLEP-seq), a long-read approach built on full-length elongating and polyadenylated RNA sequencing (FLEP-seq)^26^. Nanopore sequencing of chromatin-associated RNA gives low per-locus coverage, which is especially limiting for low-abundance transcripts such as *FLC* and *COOLAIR*. tFLEP-seq overcomes this with in-solution hybridization capture, allowing us to quantify poly(A) and non-poly(A) chromatin-associated RNA, map their 3’ ends and resolve splicing patterns across the locus throughout vernalization (Figure 1A). We enriched *FLC*, *COOLAIR* and 15 other loci^20^ from chromatin-associated RNA across a vernalization time course, achieving ∼500-fold enrichment over genome-wide background (Figure S2A). The previously described *COOLAIR* class I, II and III poly(A) isoforms were clearly detected, and the sequencing depth resolved poly(A) site clusters, termination windows and splicing efficiency in greater detail than before.

**Figure 1.**
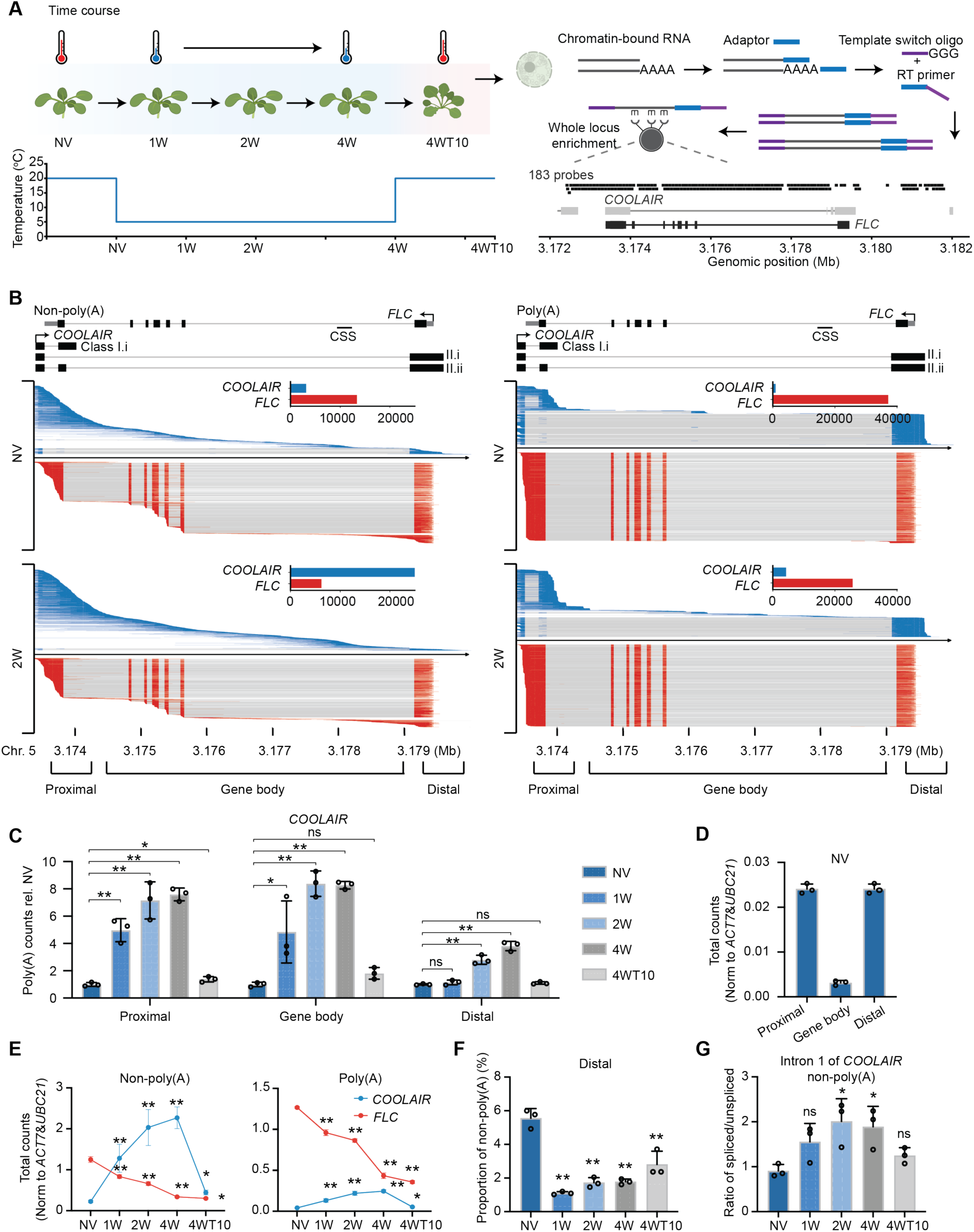
Co-transcriptional processing changes under constant cold exposure. **A.** Schematic of the tFLEP-seq experimental workflow. At each time point, chromatin-bound RNA was isolated. Full-length cDNA libraries were generated and subjected to in-solution hybridization capture targeting the *FLC*/*COOLAIR* locus and 15 additional loci. NV: non-vernalized, 1W: one week cold, 2W: 2 weeks cold, 4W: four weeks cold, 4WT10: four weeks cold and ten days post-cold. **B.** Distribution of non-poly(A) and poly(A) RNA sequencing reads across the *FLC* locus over the vernalization time course. The inset bar plot (top right) quantifies the number of reads of *COOLAIR* antisense transcripts and *FLC* sense transcripts. The black boxes and lines indicate gene models. **C.** Poly(A) transcript changes in the three windows over the vernalization time course. See methods for window definitions. Poly(A) read counts were normalized to the geometric mean of *ACT7* and *UBC21* and then normalized to the NV (non-vernalized) condition to show changes in transcript levels relative to NV. **D.** Relative abundance (normalized to the geometric mean of *ACT7* and *UBC21*) of poly(A) transcripts detected by tFLEP-seq, in different parts of *COOLAIR* locus in NV condition. **E.** Normalized read counts of *FLC* and *COOLAIR* over the vernalization time course. Reads were normalized to the geometric mean of *ACT7* and *UBC21* as done for the *FLC* strand^20^. **F.** Proportion of *COOLAIR* non-poly(A) reads with 3’ ends in the distal window calculated relative to the total *COOLAIR* non-poly(A) reads across the locus, for each timepoint. **G.** Overall ratio of spliced/unspliced reads in non-poly(A) transcripts from tFLEP-seq at *COOLAIR* strand over the time course. *n* = 3 biological replicates at all timepoints. Error bars indicate mean ± standard deviation in panels **C-G**. Statistical comparisons were performed using Tukey’s HSD (ns indicates no significance; * indicates adjusted *p* < 0.05; ** indicates adjusted *p* < 0.01).

Non-poly(A) reads in the sense direction showed that *FLC* transcription decreased gradually with increasing cold exposure. Their 3’ ends accumulated predominantly at the *FLC* transcription termination site (Figure 1B), with higher 3’ end signal over exons^27^ (Figure 1B). Despite the high read depth, we did not detect *COLDAIR* or *COLDWRAP*, two non-poly(A) lncRNAs previously described using PCR^28,29^. In the antisense direction, the non-poly(A) *COOLAIR* 3’ end profile differed markedly from that of *FLC*, with signal accumulating progressively in different regions of the locus over 4 weeks of cold, then returning to non-vernalized levels after 10 days in the warm (Figure 1E and Figure S3). This progressive increase is consistent with the gradual cold induction of CBFs^30^ and NTL8^31^, transcription factors that promote *COOLAIR* transcription. Significant accumulation of non-poly(A) 3’ ends at the proximal *COOLAIR* poly(A) site and at the CSS supports Pol II stalling at these sites, consistent with the NET-seq data (Figure S3). Moreover, *COOLAIR* non-poly(A) fold changes exceeded those in the poly(A) fraction and in NET-seq (Figure 1E). Whereas NET-seq reports Pol II occupancy, tFLEP-seq measures chromatin-associated RNA more broadly; the larger non-poly(A) response therefore suggests differential chromatin retention of *COOLAIR* transcripts.

We next analysed *COOLAIR* co-transcriptional changes in three windows across the locus: proximal, gene body and distal. Over the constant-cold time course, *COOLAIR* transcripts accumulated in all three windows, although the early responses and polyadenylation patterns differed between them (Figure 1B-D). Within one week of cold, poly(A) *COOLAIR* increased in the proximal and gene-body windows, whereas both poly(A) and non-poly(A) *COOLAIR* decreased in the distal window (Figure S2B and Figure 1F). This indicates cold-induced changes in Pol II processivity and termination, consistent with the NET-seq data. Two weeks of cold enhanced splicing of the first *COOLAIR* intron (Figure 1G). Rapid polyadenylation changes and slower splicing changes therefore differentially shift the proportions of *COOLAIR* isoforms known to repress *FLC*. Despite the initial decrease in total distal poly(A) *COOLAIR*, distal class II.ii *COOLAIR*, which represses *FLC* by sequestering FRIGIDA in condensates^9^, increased rapidly in both absolute abundance and relative proportion. In contrast, distally spliced class II.i *COOLAIR*, which directly influences *FLC* transcription^8^, and proximally spliced *COOLAIR*, linked to transcription-coupled repression^15^, increase more slowly.

### Co-transcriptional processing on the *COOLAIR* strand captures signatures of fluctuating temperatures

*COOLAIR*-mediated repression of *FLC* transcription is particularly important under natural field conditions, where dips below freezing super-induce *COOLAIR*^19^. Given the co-transcriptional sensitivity of *COOLAIR* in constant cold, we analysed its response to a regime fluctuating to below freezing (fluctuating strong, FS)^19^, compared with constant cold at the same mean temperature (5 °C) (Figure 2A). To determine whether FS alters *COOLAIR* co-transcriptional processing beyond its previously established transcriptional induction, we analysed chromatin-associated RNA using tFLEP-seq. This revealed higher levels of both non-poly(A) and poly(A) *COOLAIR* in the proximal and gene-body windows than under constant cold. Most notably, both non-poly(A) and poly(A) 3’ ends accumulated strongly around the CSS (Figure 2B-D and Figure S4A). Poly(A) *COOLAIR* transcripts terminating around the CSS, which we term class IV and class V, made up only ∼5-10% of chromatin-bound poly(A) *COOLAIR* in constant cold but ∼40% under FS (Figure 2E). Class IV and V transcripts were enriched in the chromatin bound fraction (Figure 2B, C) and, within the limits of read length, appeared unspliced (Figure 2B). Constant cold enhanced both *COOLAIR* expression and splicing efficiency, whereas fluctuations below freezing super-induced *COOLAIR* without further increasing splicing efficiency (Figure 2F). FS therefore induces a distinct *COOLAIR* co-transcriptional response, characterized by increased termination around the CSS without a corresponding increase in splicing efficiency, rather than simply amplifying the changes induced by constant cold. As predicted by modelling of field data^32^, FS did not enhance induction of *VIN3* (Figure S4B), which encodes the Polycomb accessory protein required for epigenetic silencing3. Consistent with previous evidence that FS does not enhance H3K27me3 accumulation compared with constant cold^3^, the additional *COOLAIR*-associated repression under FS is unlikely to act through enhanced H3K27me3 nucleation.

**Figure 2.**
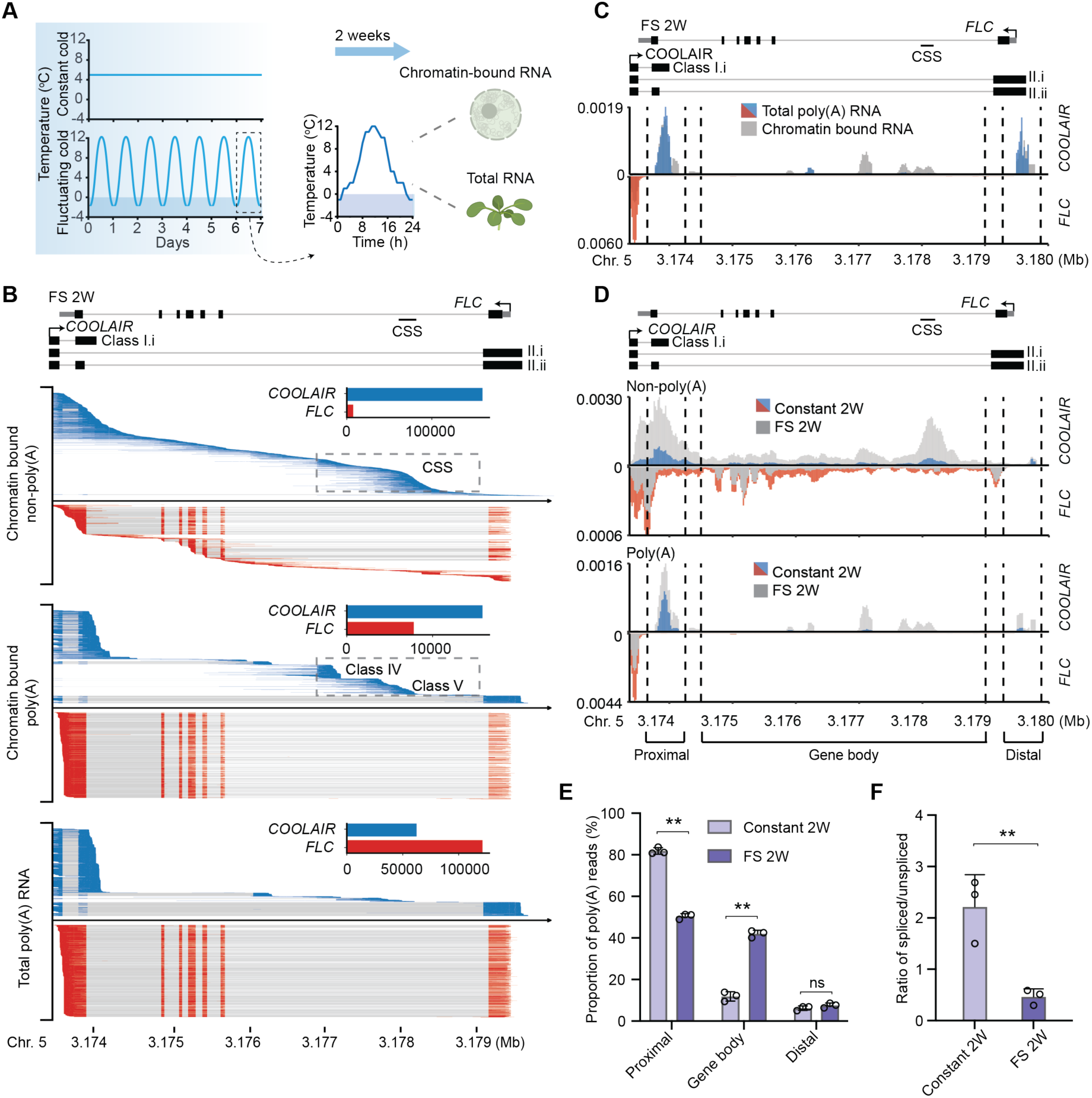
Distinct co-transcriptional changes in response to fluctuating cold. **A.** Plants were subjected to either constant cold or a fluctuating cold regime with the same mean temperature (5 °C). Following treatment, both chromatin-bound RNA and total RNA were extracted in parallel from each condition. **B.** Distribution of chromatin-bound non-poly(A), chromatin-bound poly(A) and total poly(A) RNA sequencing reads across the *FLC* locus in fluctuating strong (FS) temperature condition. The inset bar plot (top right) quantifies the number of reads of *COOLAIR* antisense transcripts and *FLC* sense transcripts. **C.** Chromatin bound non-poly(A), chromatin bound poly(A) and total poly(A) RNA 3’ end distributions across *FLC* locus in FS condition. Reads signal was normalized to the geometric mean of *ACT7* and *UBC21*. The *COOLAIR* stall site (CSS) is indicated by black bold line. **D.** Chromatin bound RNA 3’ end distributions across *FLC* locus in constant cold and FS conditions. Read signal was normalized to the geometric mean of *ACT7* and *UBC21*. **E.** Proportion of *COOLAIR* poly(A) reads with 3’ ends in the proximal, gene body and distal window calculated relative to the total *COOLAIR* poly(A) reads across the locus. See Figure 1 for window definition. *n* = 3 biological replicates at all timepoints. Statistical comparisons of the means were performed using Tukey’s HSD (ns indicates no significance; ** indicates adjusted *p* < 0.01). **F.** Ratio of spliced/unspliced reads in non-poly(A) transcripts from tFLEP-seq at *COOLAIR* strand in constant cold and FS conditions. *n* = 3 biological replicates at all timepoints. Error bars indicate mean ± standard deviation in panels **E-F**. Statistical comparison of the means was performed using a two-sided Welch’s t-test (** indicates *p* < 0.01).

### Cold-induced transcriptional changes enhance R-loop formation and *COOLAIR* clouds

The strong increase in chromatin-retained *COOLAIR* under FS prompted us to investigate the nature of this RNA. In warm conditions, *COOLAIR* transcription forms an extensive RNA-DNA hybrid over the proximal region that is associated with *FLC* repression^33^, and proximal *COOLAIR* is likely to be produced when this R-loop is resolved^34^. We used DRIPc (DNA-RNA immunoprecipitation followed by cDNA conversion) to map R-loops strand-specifically. Proximal R-loop levels increased in both constant cold and FS (Figure 3A), consistent with the increase in chromatin-associated proximal *COOLAIR*. We did not detect R-loops over the CSS, suggesting that *COOLAIR* retention at this site involves other mechanisms.

**Figure 3.**
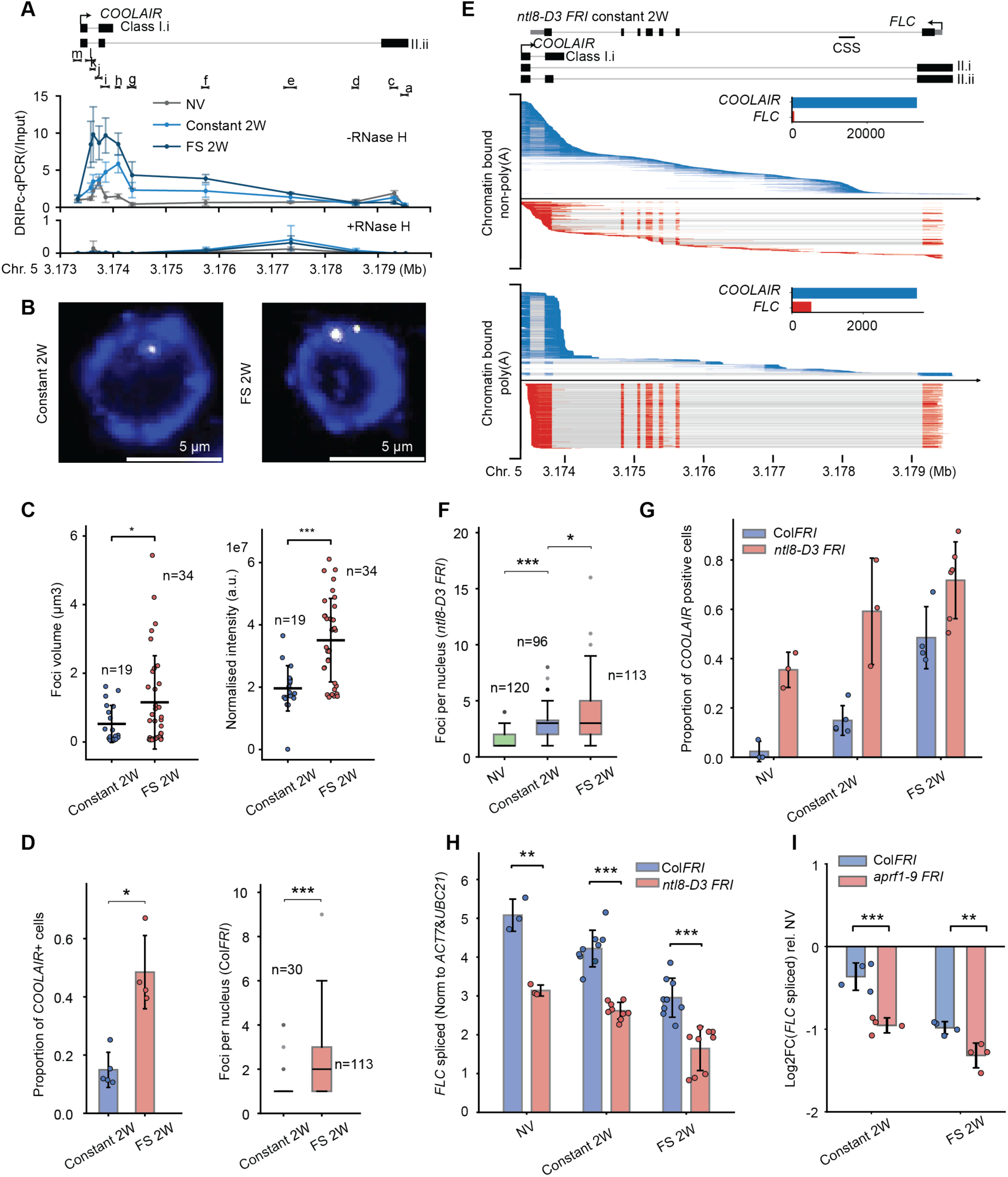
Chromatin-retained *COOLAIR* RNA induction involves complex layers of regulation. **A.** DRIPc-qPCR analyzing the R-loop over *COOLAIR* gene locus in NV, constant cold 2W and FS 2W, with and without RNase H treatment. X-axis shows the genomic position. Error bars indicate mean ± standard deviation from three independent experiments. **B.** Representative smFISH images showing spatial distribution of individual *COOLAIR* RNA foci (white) in root meristem cells. Nuclear staining from DAPI is shown in blue. Central regions with reduced DAPI staining are indicative of nucleoli. Scale bars = 5 µm. **C.** *COOLAIR* foci quantification in Col*FRI* by smFISH. Dots indicate foci volume (μm^3^) (left) and normalized intensity (a.u.) (right). Constant cold samples (blue), FS samples (red). Error bars indicate mean ± standard deviation. Number of cells in each sample indicated. In both cases, a one-sided Welch’s t-test was used to compare the means (* indicates *p* < 0.05; *** indicates *p* < 0.001). **D.** *COOLAIR* foci in cold conditions quantified in Col*FRI.* Left panel: Proportion of cells showing *COOLAIR* foci, measured by Whole-mount smFISH in root tips (see Methods). Right panel: Number of *COOLAIR* foci per nucleus measured Whole-mount smFISH in root tips. n indicates the number of cells included in the quantification. Constant cold condition: blue; FS condition: red. In both cases, a one-sided Welch’s t-test was used to compare the means (* indicates *p* < 0.05; *** indicates *p* < 0.001). **E.** Distribution of chromatin-bound non-poly(A) and chromatin-bound poly(A) reads of *ntl8-D3 FRI* across the *FLC* locus in constant cold condition. The inset bar plot (top right) quantifies the number of reads of *COOLAIR* antisense transcripts and *FLC* sense transcripts. **F.** *COOLAIR* foci per nucleus quantified in *ntl8-D3* by Whole-mount smFISH in root tips. Non-vernalized: green; Constant cold condition: blue; FS condition: red. One-sided Welch’s t-tests were used to compare the means and *P*-values were adjusted using a Bonferroni correction (* indicates adjusted *p* < 0.05; *** indicates adjusted *p* < 0.001). **G.** Comparing the proportion of *COOLAIR* positive cells in Col*FRI* and *ntl8-D3* measured by Whole-mount smFISH in root tips under different conditions. Col*FRI*: blue; *ntl8-D3*: red. **H.** *FLC* transcriptional output (normalized to the geometric mean of *PP2A* and *UBC*) between FS and constant cold conditions in Col*FRI* and *ntl8*-*D3 FRI* mutant, using a two-sided Student’s t-test (** indicates *p* < 0.01; *** indicates *p* < 0.001). **I.** Comparison of changes in *FLC* transcriptional output measured by qPCR between FS and constant cold conditions in Col*FRI* and *aprf1-9 FRI* mutant. In each condition, log_2_FC for *FLC* spliced transcript in Col*FRI* and *aprf1-9 FRI* were compared using a two-sided Student’s t-test (** indicates *p* < 0.01; *** indicates *p* < 0.001).

The increase in chromatin-retained *COOLAIR* under FS detected by tFLEP-seq reflects an average across many cells and *FLC* copies. We therefore used single-molecule FISH (smFISH) to determine whether this increase reflected greater accumulation at individual loci, a greater fraction of cells with detectable *COOLAIR* foci, or both. Previous smFISH showed that in constant cold *COOLAIR* forms large foci at the locus^18^, termed ‘clouds’ by analogy with the *Xist* lncRNA that coats the inactive X chromosome^35^. Here we used *COOLAIR* intronic probes that also cover the CSS. As previously reported^24^, *COOLAIR* clouds were present in a small fraction of cells under constant cold. Under FS, clouds increased in both size and intensity, indicating greater *COOLAIR* accumulation at loci with detectable clouds (Figure 3B, C). The number of foci per nucleus and the fraction of cells with foci also increased (Figure 3D), consistent with more frequent *COOLAIR* transcription. Clouds form for *COOLAIR* but not *FLC*^18^, and tFLEP-seq detected more non-poly(A) *COOLAIR* than non-poly(A) *FLC* (Figure 1E). Together, these observations suggest that the clouds comprise chromatin-retained non-poly(A) *COOLAIR* at each allele. Thus, constant cold and FS alter *COOLAIR* co-transcriptional regulation (Pol II processivity, termination and transcript release) through distinct modes at different sites within the locus.

### Genetic perturbations distinguish contributions of *COOLAIR* transcription and processing to *FLC* repression

We next asked which *COOLAIR* co-transcriptional changes are functionally important, using mutants in which we had previously established a causal link between altered *COOLAIR* regulation and *FLC* expression. NTL8 is a transcription factor that binds directly to the *COOLAIR* 5’ region^19^. The dominant *ntl8-D3 FRI* allele ectopically expresses high levels of *COOLAIR*, resulting in *FLC* hyper-repression^19^. tFLEP-seq confirmed high *COOLAIR* expression in *ntl8-D3 FRI*, which increased further after 2 weeks of constant cold (Figure 3E), accompanied by further *FLC* repression (Figure 3H). Non-poly(A) *COOLAIR* increased significantly over the proximal region and parts of the gene body but importantly not at the CSS (Figure 3E, Figure S4C). While poly(A) *COOLAIR* significantly increased over the proximal region, the proportion of class IV and V poly(A) *COOLAIR* did not increase in *ntl8-D3 FRI* (Figure S4D). The increased proportion of class IV and V polyadenylation under FS is therefore not simply a consequence of increased *COOLAIR* transcription. *COOLAIR* clouds, which report gene-body chromatin-retained transcripts, were enhanced in *ntl8-D3 FRI* in the warm and further enhanced by constant cold and FS (Figure 3F, G), with *FLC* repression strongest under FS (Figure 3H). Together, these data suggest that elevated *COOLAIR* transcription is sufficient to enhance *FLC* repression, whereas additional FS-induced co-transcriptional changes, including those at the CSS, strengthen it further.

We had previously shown that a hypomorphic *smu1 FRI* mutant disrupts cold-enhanced splicing of the proximal intron and *COOLAIR* proximal polyadenylation^12^. This effect was phenocopied by cis mutation of the 3’ splice acceptor site, causally linking these *COOLAIR* processing changes to *FLC* repression^12^. tFLEP-seq analysis of *smu1 FRI* after 2 weeks of constant cold confirmed reduced proximal intron splicing and proximal polyadenylation relative to Col*FRI* (Figure S5A-E), accompanied by impaired *FLC* repression (Figure S5B). By contrast, under FS, *smu1 FRI* still showed reduced proximal splicing relative to Col*FRI*, to a similar extent as in constant cold (Figure S5F), but proximal polyadenylation was not reduced relative to Col*FRI*, and *FLC* repression was retained (Figure S5G, I). Under FS, non-poly(A) *COOLAIR* and poly(A) classes IV and V also differed little between *smu1 FRI* and Col*FRI* (Figure S5H-J). These results therefore show that reduced proximal splicing alone is insufficient to fully reduce *FLC* repression in FS, and that its functional effect depends on both the accompanying polyadenylation pattern and the level of *COOLAIR* transcription.

We then examined *aprf1-9 FRI*, which in the warm increases gene-body polyadenylation and reduces distal polyadenylation of *COOLAIR*^15^, mimicking the cold-induced shift (Figure 1 and 2). APRF1 interacts with components of the phosphatase module of the CPF-like 3’ processing complex that regulates *COOLAIR* processing^15^. tFLEP-seq of *aprf1-9 FRI* in the warm revealed a polyadenylation shift consistent with previous data (Figure S6A). However, this shift does not produce detectable *FLC* repression in the warm^15^, likely because *COOLAIR* transcription remains low. To probe the link between *COOLAIR* polyadenylation and *FLC* repression, we asked whether the shift in *aprf1-9 FRI* persists when cold induces *COOLAIR*, and whether it enhances cold-induced *FLC* repression. The polyadenylation shift persisted in constant cold (Figure S6A), and *FLC* repression was enhanced. Repression in *aprf1-9 FRI* under constant cold was comparable to that in wild type under FS (Figure 3I). This suggests that increased *COOLAIR* transcription in the cold strengthens the repressive effect of the polyadenylation shift by increasing the frequency of processing and termination events at the favoured sites. Together with the temperature-dependent *COOLAIR* processing profiles, these results functionally implicate polyadenylation and termination events associated with the CPF phosphatase module, which occur predominantly in the proximal region in constant cold and extend into the gene body under regimes fluctuating below freezing.

Together, these perturbations show that *COOLAIR*-mediated *FLC* repression depends on both the level of *COOLAIR* transcription and the accompanying pattern of co-transcriptional processing. The functional consequences of altering a particular processing step therefore depend on the broader transcriptional and processing response to the temperature regime.

### *COOLAIR* co-transcriptional processing is linked to local chromatin interactions

Pol II transcription can shape fine-scale chromatin organization at individual genes^36,37^, including in plants^38,39^, where such interactions change in response to cold^40^. Chromosome conformation capture (3C) previously showed that cold disrupts a gene loop linking the *FLC* transcription start site (TSS) and transcription termination site (TTS), which is also close to the *COOLAIR* TSS^17^, and that this loop is affected by *COOLAIR* transcription^3^. We therefore asked whether changes in local chromatin interactions could link *COOLAIR* co-transcriptional changes, and the associated redistribution of Pol II and chromatin-retained RNA, to *FLC* repression. We performed Micro Capture-C^41^ with a whole locus enrichment step (Methods), enabling single-gene-resolution mapping of chromatin interactions across a 10-kb region around *FLC* in warm, constant cold and FS conditions (Figure 4A, Figure S6B). In the warm, the normalized contact map (Figure 4B, upper triangles) revealed two major local interaction domains. One spans the *FLC* gene from TSS to TTS. The other, upstream, covers the *FLC* promoter and extends to the TTS of the neighbouring gene *UPSTREAM OF FLC* (*UFC*; AT5G10150). Micro-C-XL previously detected both domains in Col-0, where *FLC* is epigenetically silenced^39^, as subdomains of a larger 20-kb domain in which the *FLC* gene body contacts its upstream region; in those data, the promoter domain was separate from *UFC*. Both cold treatments disrupted the *FLC* gene-domain boundaries through the emergence of new interactions (Figure 4B and Figure S6C). To account for distance-dependent decay and highlight local interaction patterns, we computed observed/expected (O/E) matrices for each condition (Figure 4B, lower triangles). These showed enriched interactions within the *FLC* gene domain, consistent with the gene loop previously detected by 3C^17^. Both constant cold and FS reduced interactions within the gene domain, correlating with significantly reduced *FLC* transcription (Figure 4B and Figure S6D).

**Figure 4.**
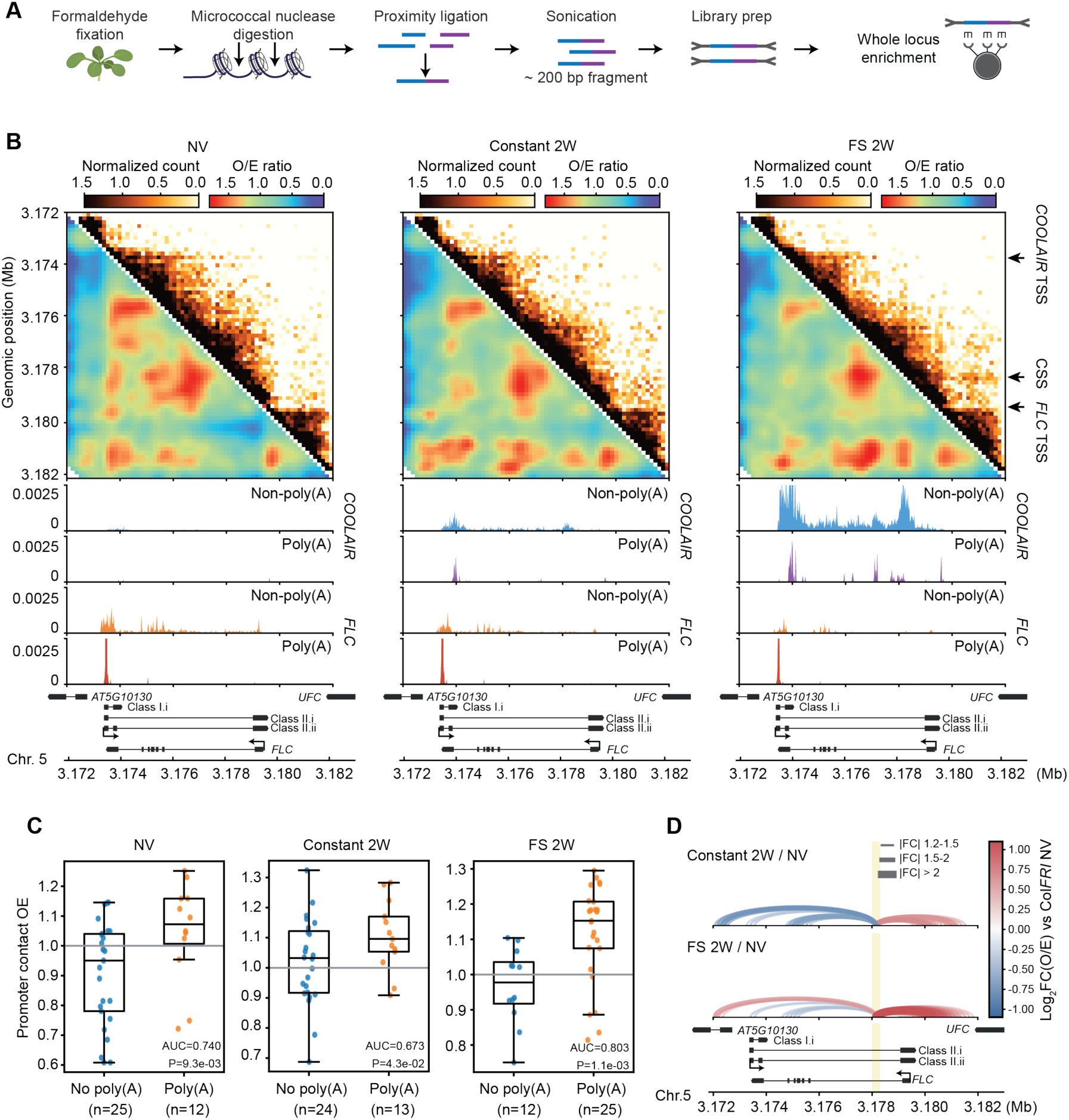
Temperature responsive changes in *COOLAIR* co-transcriptional processing are linked to chromatin changes affecting *FLC* transcription A. Chromatin was digested with Micrococcal nuclease, followed by proximity ligation. The resulting library was subjected to in-solution hybridization capture, enabling high-resolution mapping of chromatin contacts across a 10-kb region at single-gene resolution. **B.** Micro Capture-C contact maps of the *FLC* locus (Chr5 3.172-3.182 Mb) under three conditions: NV, constant cold 2W, and FS 2W. For each condition, the upper-right triangle displays normalized interaction counts and the lower-left triangle displays O/E ratios, with color scales shown above each map. Gene annotations are shown below each map. Two boundaries (*COOLAIR* TSS and *FLC* TSS) and *COOLAIR* stalling site (CSS) positions are indicated by arrows on the right. The 3’ end positional distributions of non-poly(A) and poly(A) *COOLAIR* transcripts are also shown as coverage tracks aligned to the contact map coordinates. Read signal was normalized to the geometric mean of *ACT7* and *UBC21*. **C.** Promoter-contact observed/expected (OE) values for regions with and without detected *COOLAIR* polyadenylation in NV, constant cold 2W and FS 2W conditions. Points represent individual 200-bp bins; boxes show the median and interquartile range, with whiskers extending to 1.5 times the interquartile range. The grey line indicates OE = 1. AUC (area under the receiver operating characteristic curve) gives the probability that a randomly chosen poly(A)-positive region has higher OE than a randomly chosen poly(A)-negative region; *P* values were obtained by randomly reassigning the observed number of poly(A)-positive labels among the 200-bp bins and recalculating the AUC. **D.** Micro-Capture-C contact comparisons between the CSS and the 10-kb region under the indicated conditions. Interaction fold change was calculated from observed/expected (OE) matrices normalized for genomic distance. Red and blue indicate increase and decrease in interactions, respectively, and the line width represents the magnitude of the fold change.

The O/E matrices also revealed prominent contacts between the promoter domain and *FLC* gene regions enriched for *COOLAIR* poly(A) sites; promoter-contact O/E values were consistently elevated at regions of preferential *COOLAIR* polyadenylation (Figure 4C and Figure S6E). In the warm, these contacts were concentrated over the proximal *COOLAIR* poly(A) region. Constant cold enhanced contacts with both proximal and gene-body regions, paralleling the cold-induced shift in *COOLAIR* polyadenylation. This correspondence was strongest under FS, when a prominent promoter contact emerged over the CSS (Figure 4D). This is the site where Pol II accumulated in the cold and where, under FS, non-poly(A) and class V poly(A) *COOLAIR* increased. Together, these observations support a model in which the spatial pattern of *COOLAIR* 3’ processing and Pol II termination helps determine where promoter-gene contacts form. Whereas termination-associated gene looping has been described previously^42–45^, our data link this mechanism to environmentally regulated antisense transcription and to dynamic repositioning of contacts within a single gene.

To test this association genetically, we examined three independent perturbations. The first was a *COOLAIR* mutant (*FLCΔCOOLAIR*) that disrupts cold induction of *COOLAIR* transcription^9,19,46^. The second was *ntl8-D3 FRI*, which constitutively increases *COOLAIR* transcription, and the third was *aprf1-9 FRI*, which enhances gene-body *COOLAIR* polyadenylation. For each, we quantified the proportion of all observed contacts that occurred between the promoter domain and the *FLC* gene. In the *COOLAIR* mutant, this proportion remained significantly lower than in Col*FRI*, even under FS (Figure 5A, C), consistent with the negligible poly(A) *COOLAIR* detected over the locus (Figure S7A). *ntl8-D3 FRI* did not increase the promoter-gene contact proportion relative to Col*FRI* in either warm or cold, despite significantly higher *COOLAIR* transcription (Figure 5C and Figure S7B, C). Increased *COOLAIR* transcription alone is therefore insufficient to enhance promoter-gene interactions. By contrast, *aprf1-9 FRI* increased the promoter-gene contact proportion in both warm and cold, corresponding to its increased gene-body *COOLAIR* polyadenylation (Figure 5A, B, D). In *aprf1-9 FRI*, elevated promoter contacts again coincided with regions of preferential *COOLAIR* polyadenylation in both conditions (Figure 5E). This shows that genetically redistributing *COOLAIR* 3’ processing also redistributes promoter contacts. Together, these perturbations indicate that promoter contacts track the spatial pattern of *COOLAIR* 3’ processing rather than *COOLAIR* transcription level.

**Figure 5.**
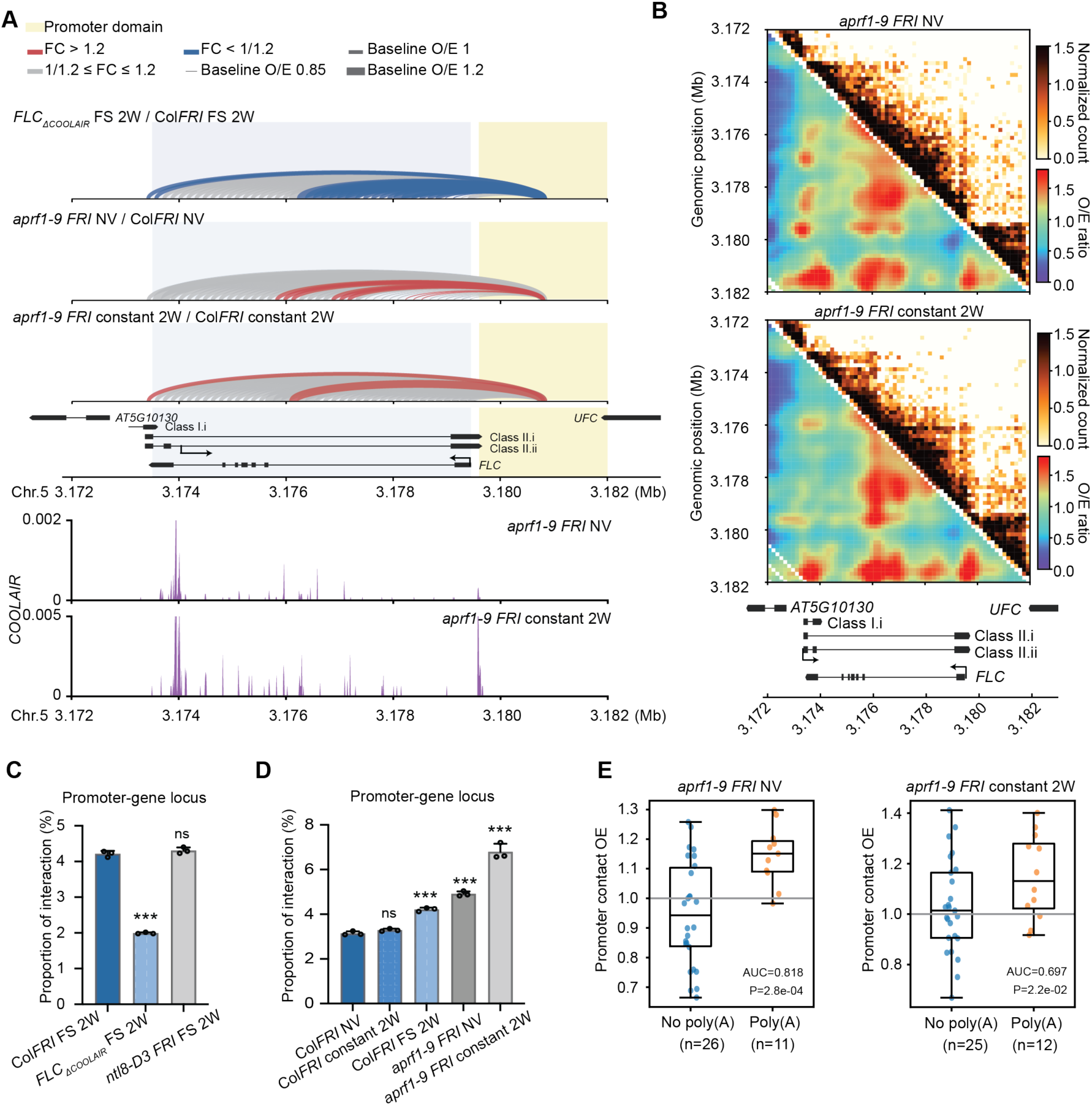
APRF-dependent co-transcriptional regulation modulates chromatin interactions at the *FLC* locus. **A.** Micro-Capture-C contact profiles between the promoter region and the *FLC* gene locus in the indicated mutants. The top panel shows the contact distributions for the different mutants. The bottom panel shows aligned poly(A) read signals for *aprf 1-9 FRI* under NV and constant 2W conditions. Read signals were normalized to the geometric mean of *ACT7* and *UBC21* reads. Arcs were plotted using the log_2_ (fold change) in O/E value. Arc colour and width represent the fold change in OE and baseline OE value in the control, respectively. **B.** Micro Capture-C contact maps of the *FLC* locus (Chr5 3.172-3.182 Mb) in *aprf1-9 FRI*. For each condition, the upper-right triangle displays normalized interaction counts and the lower-left triangle displays O/E ratios, with color scales shown above each map. Gene annotations are shown below each map. **C.** Proportion of interactions between the promoter region and the *FLC* gene locus relative to the total reads across the 10-kb region in Col*FRI*, *FLC_ΔCOOLAIR_ FRI*, and *ntl8-D3 FRI* under FS conditions. **D.** Proportion of interactions between the promoter region and the *FLC* gene locus in Col*FRI* under NV, constant 2W, and FS conditions, and in *aprf1-9 FRI* under NV and constant 2W conditions. In **C** and **D**, statistical significance was assessed using two-side Students’s t-tests. *P* values were adjusted using the Holm-Sidak approach (ns indicates no significance; * *p* < 0.05; ** *p* < 0.01; *** *p* < 0.001). **E.** Promoter-contact observed/expected (OE) values for regions with and without detected *COOLAIR* polyadenylation in NV and constant cold conditions. Points represent individual 200-bp bins; boxes show the median and interquartile range, with whiskers extending to 1.5 times the interquartile range. The grey line indicates OE = 1. AUC gives the probability that a randomly chosen poly(A)-positive region has higher OE than a randomly chosen poly(A)-negative region; *P* values were obtained by randomly reassigning the observed number of poly(A)-positive labels among the 200-bp bins and recalculating the AUC.

### Distinct temperature regimes induce different local chromatin modification patterns

We then asked how these co-transcriptional and chromatin-interaction changes relate to chromatin modifications and *FLC* expression. We previously showed that Pol II termination promotes a repressive chromatin state at *FLC* through H3K4me1 demethylation^15,16,47^, together with the activity of APRF1 and the phosphatase module of the CPF-like complex^15^. We measured H3K4me1 across the locus under FS and compared it with constant-cold time-course data^3^. In both regimes, H3K4me1 decreased over the proximal and gene-body regions (Figure 6A, B). This is consistent with increased conversion to H3K4me3 in the proximal window, associated with increased *COOLAIR* transcription (Figure S8A). It is also consistent with reduced co-transcriptional deposition of H3K4me1 in the gene body, owing to reduced sense transcription (Figure S8B). Other chromatin changes were detectable after one week of constant cold, including reduced H3 acetylation (Figure S8C) and repositioning of the +1 nucleosome^48^. In a distinct genetic context, nucleosome repositioning at *FLC* also accompanied *COOLAIR* co-transcriptional changes in *ntl8-D3* in the absence of cold (Figure S9A, B). This supports a mechanistic link between antisense co-transcriptional changes and local nucleosome dynamics, as reported in yeast^49,50^.

**Figure 6.**
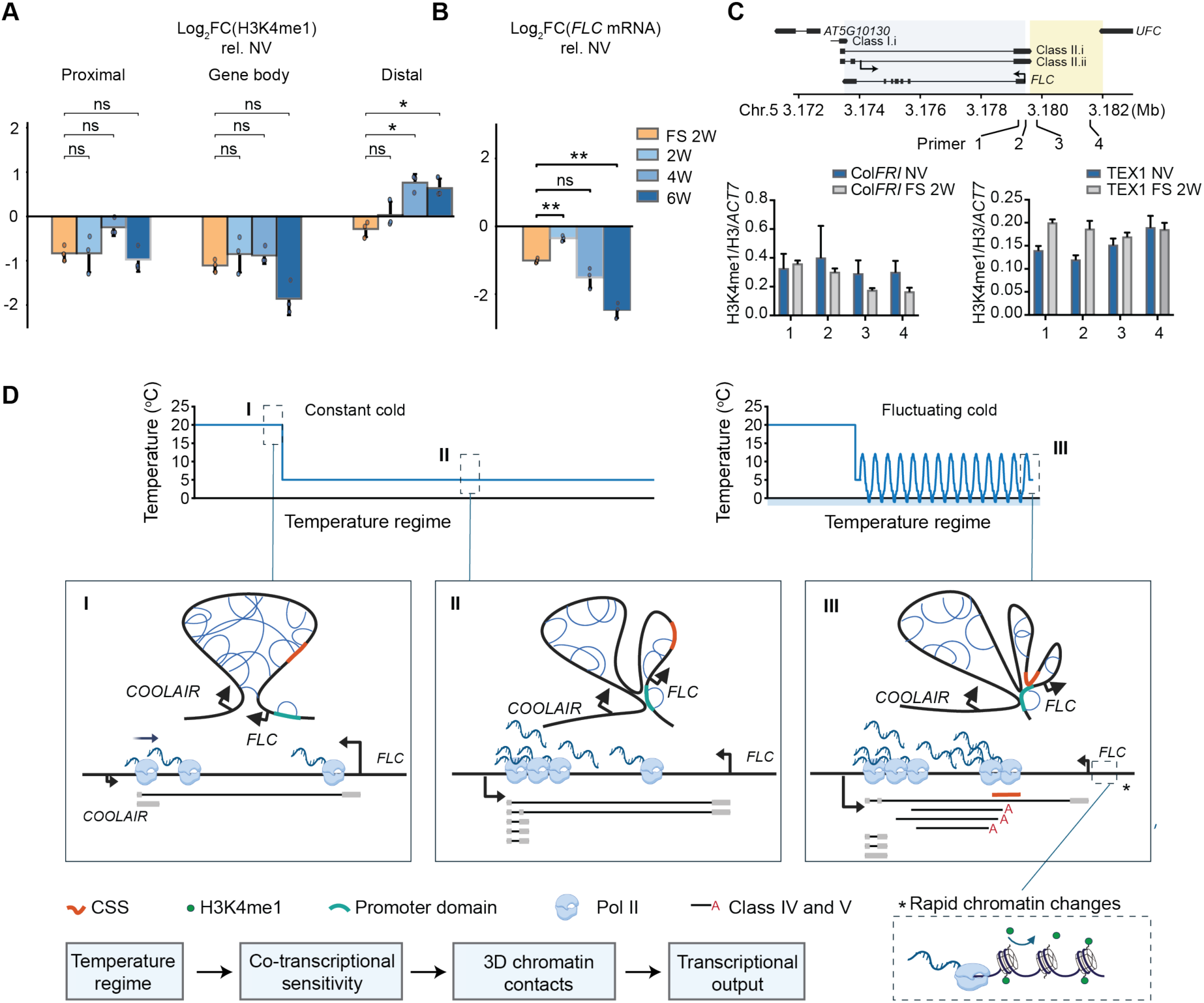
*COOLAIR* co-transcriptional processing serves as thermosensory antennae that integrate different facets of complex winter temperature profiles A. Comparison of H3K4me1 changes between FS and constant cold conditions in the three regions defined above. Constant cold data reproduced from Nielsen et al., 2024^3^.Average log_2_ fold change (Log_2_FC) relative to NV was computed over multiple primers covering each region, after normalizing to H3 and the control locus *ACT7*. For each region log_2_FC at different constant cold timepoints were compared to log_2_FC under FS condition, using a two-sided Student’s t-test. *P* values were adjusted using the Holm-Sidak approach (ns indicates no significance; * indicates adjusted *p* < 0.05). **B.** Comparison of changes in *FLC* transcriptional output between FS and constant cold conditions. FS and constant cold data reproduced from Nielsen et al., 2024^3^. Log_2_FC at different constant cold timepoints were compared to log_2_FC under FS condition, using a two-sided Student’s t-test. *P* values were adjusted using the Holm-Sidak approach (ns indicates no significance; * indicates adjusted *p* < 0.05; ** indicates adjusted *p* < 0.01). **C.** H3K4me1 changes across the *FLC* locus measured by ChIP-qPCR under FS conditions, in the *COOLAIR* defective mutant line TEX1.0. Magnified view of the profile over the *FLC* promoter/upstream region in Col*FRI* (Left) and TEX1.0 (Right). **D.** Schematic model of the co-transcriptional regulation of *COOLAIR* transcription. *COOLAIR*-mediated *FLC* repression responds to temperature changes over multiple timescales. Different winter temperature regimes affect not only transcriptional initiation but also RNA Pol II progression and RNA processing during *COOLAIR* transcription. Early during cold exposure, increased RNA Pol II stalling within the *COOLAIR* gene body limits polymerase progression towards its 3’ end, near the *FLC* transcription start site. With prolonged cold exposure, proximal *COOLAIR* transcripts progressively accumulate. Fluctuations below freezing (FS) further enhance RNA Pol II stalling and promote premature termination within the gene body, generating both polyadenylated and non-polyadenylated *COOLAIR* 3’ ends. Regions of preferential *COOLAIR* 3’ processing are enriched for contacts with the *FLC* promoter, and genetic perturbations of processing alter the distribution of these contacts. Under FS, enhanced promoter-gene-body contacts coincide with reduced H3K4me1 and *FLC* transcription.

A clear difference emerged over the distal *COOLAIR*/*FLC* promoter region, immediately downstream of the CSS. After prolonged constant cold (4 and 6 weeks), H3K4me1^3^ (Figure 6A, B) and H3K4me2^51^ (Figure S8A) increased in the distal region. This is consistent with reduced conversion of H3K4me1 to H3K4me2, and of H3K4me2 to H3K4me3, as initiation of *FLC* sense transcription declines. Under FS, however, although *FLC* repression was similar in magnitude to that after 4 weeks of constant cold, distal H3K4me1 did not increase but instead decreased slightly (Figure 6A, B). These changes were disrupted in the *COOLAIR*-defective TEX1.0 line (Figure 6C), indicating that they depend on *COOLAIR* and supporting a role for antisense co-transcriptional processing in shaping local chromatin state. The loss of H3K4me1 over the distal *COOLAIR* region and *FLC* 5’ end under FS suggests active demethylation, linked to enhanced *COOLAIR* termination at the CSS and the associated changes in local chromatin topology.

## Discussion

Understanding how organisms integrate complex, fluctuating environmental signals is a fundamental problem in biology. We addressed this question by studying the cold-induced repression of the *Arabidopsis* floral repressor *FLC*. Our key finding is that distinct *COOLAIR* co-transcriptional processes act as multiple temperature antennae operating over different timescales, providing a mechanism to integrate different features of winter temperature. Beyond changes in splicing and polyadenylation, temperature strongly affected Pol II stalling and processivity, and the retention of both non-poly(A) and poly(A) transcripts on chromatin. Genetic perturbations further show that *COOLAIR*-mediated *FLC* repression depends on both the level of *COOLAIR* transcription and the accompanying pattern of co-transcriptional processing. The greater sensitivity of *COOLAIR* transcription than *FLC* sense transcription to temperature fluctuations may reflect differences in the factors loaded onto Pol II at initiation^52^.

These changes in Pol II dynamics and *COOLAIR* processing are associated with altered local chromatin interactions. Genetic perturbations suggest that the genomic distribution of *FLC* promoter contacts is associated with the locations of preferential *COOLAIR* 3’ processing. Increased *COOLAIR* transcription does not necessarily increase the proportion of promoter–gene contacts, while contacts can be enriched even when *COOLAIR* transcription remains low, as in warm *aprf1-9*. In that genotype, contact enrichment is not accompanied by detectable *FLC* repression, indicating that increased contacts alone are insufficient for repression. A possible physical basis for these processing-associated contacts is interactions between termination-associated Pol II complexes^53^, with Pol II terminating at the CSS interacting with Pol II terminating across the *FLC* promoter domain. Consistent with this, FS generates non-poly(A) *COOLAIR* 3’ ends across the promoter domain, including near the *UFC* TTS (Figure S6F). The stronger contacts under FS than in constant cold coincide with the pronounced redistribution of *COOLAIR* termination towards the CSS. Repeated temperature changes may also favour more transient transcriptional states than those occurring during prolonged constant cold, potentially contributing to the stronger contacts under FS. These altered interactions could in turn influence the recruitment or activity of chromatin-modifying factors at *FLC*.

The distinct *COOLAIR* responses to constant cold and FS are accompanied by differences in local chromatin modifications. The contrasting distal H3K4me1 responses despite comparable *FLC* repression suggest that similar transcriptional outcomes can accompany different local chromatin states. Enhanced *COOLAIR* termination near the CSS may contribute to H3K4me1 demethylation under FS, consistent with previous links between Pol II termination and demethylation^15^. H3K4me1 demethylation is one possible component of a broader chromatin response that may also involve other histone modifications and changes in chromatin organization. Such flexibility in temperature-dependent *COOLAIR* regulation enables robust *FLC* repression across winters with different temperature profiles (Figure 6D). Related antisense co-transcriptional responses are likely to also occur at other environmentally responsive genes. Genome-wide 3’ mRNA sequencing (Quant-Seq) revealed that fluctuations below freezing were associated with changed polyadenylation patterns for cold-induced antisense transcription across hundreds of genes (Figure S9C). Similar changes were also observed for protein-coding gene transcription genome wide (Figure S9D, E). Co-transcriptional plasticity may therefore be a widespread mechanism for temporally integrating naturally fluctuating signals to achieve robust gene regulation.

## Data availability

Raw sequencing data generated in this study is available under the SRA Bioproject accession number PRJNA1558601. The NET-seq data used in this study is from a previous study^17^, available under the SRA Bioproject accession number PRJNA1144665. Images generated for the smFISH analysis are available at the BioImage Archive under accession number S-BIAD3599.

## Code availability

Code used for the analysis and visualization in this study is available via GitHub (https://github.com/DingLab-RNAstructure/COOLAIR_co_transcription_analysis).

## Methods

### Plant material and growth conditions

All mutant and transgenic lines were in the *FRI^sf2^* background. *ntl8-D3 FRI^1^, aprf1-9 FRI*^2^, *smu1-6 FRI*^3^ (*smu1-7 FRI*) and *FLC_ΔCOOLAIR_* were described previously^1-4^. Seeds were surface-sterilized and sown on Murashige and Skoog (MS) agar plates without glucose or sucrose (unless otherwise stated) and kept at 4 °C in the dark for 3 days. Non-vernalized (NV) seedlings were cultivated under warm conditions (16 h light/8 h dark at a constant 20 °C) for 10 days, unless otherwise specified. For vernalization treatment, seedlings were initially grown under the same warm conditions for 10 days and subsequently moved to cold conditions (8 h light/16 h dark at a constant 5 °C) for extended periods, such as 2 or 4 weeks. For cold-to-warm transfer experiments, after 4 weeks of cold treatment, seedlings were moved to warm conditions (16 h light, 8 h darkness with constant 20 °C) for 10 days. Three biological replicate samples were taken at 9:00 after treatment.

Plants for fluctuating freezing experiments were generally grown in growth conditions as described previously^1^. Seeds were sown onto soil and kept at 4 °C in the dark for 3 days. NV plants were cultivated under warm conditions (16 h light/8 h dark at a constant 20 °C) for 10 days before plants were then moved to 8h light/16h dark growth chambers. For vernalization, plants were treated with two different temperature regimes: constant 5°C and daily fluctuating 5°C (−1°C to 12°C). Three biological replicate samples were taken at 9:00 after 2 weeks exposure to vernalization.

### RNA expression analysis

Total RNA was isolated using the previously described hot phenol protocol^5^. Genomic DNA was removed by treatment with TURBO DNase (Invitrogen) according to the manufacturer’s instructions. First-strand cDNA was generated using SuperScript IV reverse transcriptase (Invitrogen). Gene-specific primers were applied for reverse transcription of *COOLAIR* and *FLC*. Quantitative PCR (qPCR) was conducted with SYBR Green I Master (Roche) on a LightCycler 480 system (Roche). Ct values were normalized to the geometric mean of *UBIQUITIN CARRIER PROTEIN 1* (*UBC*) and *SERINE/THREONINE PROTEIN PHOSPHATASE 2A* (*PP2A*). Primer sequences are provided in Supplementary Table 1.

### tFLEP-seq library preparation and sequencing

Chromatin bound RNA extraction was performed following the protocol described^6^ with minor modifications. 2.5 g 10-day-old seedlings were harvested and grounded into a fine powder. Suspend powder in 25 mL of Honda buffer (1.25% (wt/vol) Ficoll, 2.5% (wt/vol) Dextran T40, 0.44 M sucrose, 20 mM HEPES (pH 7.4), 10 mM MgCl_2_, 0.5% (vol/vol) Triton X-100, 1 mM DTT, 1 × protease inhibitor cocktail (Roche), and 100 ng/μL tRNA), filtered through two layers of Miracloth, and centrifuged at 4 °C, 2000 × *g* for 10 min. The nuclei pellets were resuspended and washed in Honda buffer three times, or until the pellet appears colorless. The supernatant was removed thoroughly and discarded. The nuclei pellets were weighed, and one volume of nuclei resuspension buffer (25 mM Tris-HCl (pH 7.5), 50% (vol/vol) glycerol, 0.5 mM EDTA (pH 8.0), 1 mM DTT, 100 mM NaCl, 0.4 U/μL RNaseOUT and 100 ng/μL tRNA) was added to each pellet according to the weight of the pellet (e.g., 100 mg = 100 μL). Two volumes of washing buffer (25 mM Tris-HCl (pH 7.5), 300 mM NaCl, 1 M urea, 0.5 mM EDTA (pH 8.0), 1 mM DTT, 1% (vol/vol) Tween 20, 0.4 U/μL RNaseOUT and 200 ng/μL tRNA) was added, and the pellets were washed by pipetting up and down seven to ten times. The samples were centrifuged at 4 °C, 8,000 × *g*, for 1 min, the supernatant was removed and the pellets were resuspended with one volume of nuclei resuspension buffer. One volume of washing buffer was added, and the pellets were washed by pipetting up and down seven to ten times. The samples were centrifuged at 4 °C, 8,000 × *g*, for 1 min, the supernatant was removed and add 1 ml TRIzol to extract RNA.

tFLEP-seq was performed following the protocol described^6^ with modifications. rRNA depletion was performed using riboPOOL probe (siTOOLs Biotech). The RNA was cleaned and concentrated using the ZYMO RNA Clean & Concentrator-5 kit. 1 μL of 50 pmol 3’ adapter (Universal miRNA cloning linker, NEB, S1315S) was added to the rRNA-depleted RNA, followed by ligation using T4 RNA Ligase 2, truncated K227Q (NEB, M0373L). The RNA was cleaned and concentrated. Reverse transcription was carried out using a 3’ cDNA RT primer with the following sequence: 5’-phos/ACTTGCCTGTCGCTCTATCTTC-index-ATTGATGGTGCCTACAG-3’. After first-strand cDNA synthesis, PCR amplification was performed using the universal primer pair (Supplementary Table 2). Target enrichment was conducted according to the manufacturer’s instructions using the myBaits Custom DNA Targeted Sequencing Kits. The sequence of all baits is listed in Supplementary Table 3. Enriched libraries were subsequently prepared for Nanopore sequencing. Library construction was performed using the Ligation Sequencing Kit V14 for non-barcoded samples, or in combination with the Native Barcoding Kit 24 V14 when barcoding was required. Sequencing was carried out on R10 flow cell (Oxford Nanopore Technologies, FLO-MIN114) on a Minion Mk1B or PromethION 2 Solo sequencer (Oxford Nanopore Technologies). Biological replicates and correlation analysis of replicates for all samples are included in the supplementary files.

### DRIPc-qPCR

DRIPc-qPCR was performed following the protocol described^7^ with modifications. 2.5 g 10-day-old seedlings were harvested and grounded into a fine powder. The powder was suspended in 25 mL of Honda buffer, filtered through two layers of Miracloth, and centrifuged at 3500 × *g* for 15 min. According to the weight of the nuclei pellets, 5 volumes of nuclear lysis buffer (50 mM Tris-HCl (pH 7.5), 10 mM EDTA) were added into each sample. 3.5 µL of Proteinase K (20 mg/ml, AM2546) were then added per 100µL of sample from step 1, add SDS to a final concentration of 0.5% from 10% (w/v) SDS solution. The samples were incubated overnight at 55°C with gentle rotation. Equal volume of phenol/chloroform was added to each sample, mixed gently and centrifuged at 13,000 × g for 10 min. The supernatant was transferred into a new tube, and the samples were precipitated with 1 volume of Isopropanol and 1/10 volume of NaOAc. The tubes were gently inverted until DNA began to precipitate. Centrifugation was not used. DNA was spooled using pipette tips. The pellets were washed once with 70% EtOH and dried. The DNA pellets were then dissolved in 200 µL H_2_O and mixed gently until absolutely dissolved. 4 μg DNA were then used for restriction enzyme (HindIII, NdeI, XbaI) digestion overnight, followed by phenol/chloroform extraction and DNA precipitation with isopropanol. After restriction enzyme digestion and purification, aliquots of the products were treated with RNase III (NEB, M0245S) or RNase H (NEB, M0297S), followed by phenol-chloroform extraction and precipitation. 50 μL DNA (∼2 μg DNA) was diluted in ChIP dilution buffer (16.7 mM Tris-HCl (pH 7.5), 1.2 mM EDTA, 0.1% Triton X-100, 167 mM NaCl) to 495 μL, supplemented with protease inhibitors, and incubated with 5 μg S9.6 antibody (Kerafast, ENH001). A 5 μL aliquot was collected as input control and stored at 4 °C. The remaining sample was incubated overnight at 4 °C on a rotating platform. 50 μL Protein G agarose beads were then added and the samples were incubated for a further 2-3 h at 4 °C with rotation. Beads were collected by centrifugation (1000 × g, 1 min, 4 °C), washed five times with ChIP dilution buffer (5 min each), and then washed twice with TE buffer (10 mM Tris-HCl (pH 7.5), 1 mM EDTA, 0.1% Triton X-100) at room temperature. DNA-RNA hybrids were eluted by adding 200 μL Elution buffer (10 mM Tris-HCl (pH 7.5), 2 mM EDTA, 0.2% SDS, 100 ng/mL tRNA) and 5 μL Proteinase K to each sample, including input controls, followed by incubation at 55 °C for 2 h. Beads were pelleted, and the supernatant was transferred to a new tube and mixed with 300 μL H_2_O, followed by phenol/chloroform extraction and DNA precipitation with isopropanol, pelleted DNA was resuspended in 40 μL H_2_O.

DNA/RNA hybrids (excluding input samples) were denatured at 95 °C for 2 min and immediately left on ice for at least 2 min. DNA was then removed using the TURBO DNA-free kit according to the manufacturer’s instructions. RNA was precipitated by adding 1 μL glycoblue, 1/10 NaOAc, and 1 volume isopropanol, followed by overnight incubation at −20 °C. Samples were centrifuged at 12,000 × g for 15 min at 4 °C, and the RNA was washed once with 70% ethanol and centrifuged again at 12,000 × g for 5 min at 4 °C. After air-drying, RNA was resuspended in 40 μL H_2_O. A total of 10 μL RNA was used for reverse transcription reactions, which were performed separately using pooled forward primers or pooled reverse primers (Supplementary Table 4). COPIA and XCRR reverse primers were included as endogenous controls. cDNA was diluted by adding 400 μL H_2_O. For input samples, 10 μL was diluted with 410 μL H_2_O. The cDNA and input were diluted and analyzed by qPCR via LightCycler480 II (Roche).

### ChIP-qPCR

Chromatin immunoprecipitation (ChIP)-qPCR was performed as described previously^8,9^. ChIP was performed with antibodies: α-H3 (Abcam, ab1791), α-H3K4me1 (Abcam, ab8895), α-H3K4me2 (Millipore, 07-030), α-H3K4me3 (Millipore, 04-745), α-H3K9K14ac (Merck, #06-599), α-H3K27ac (Abcam, ab4729), α-H3K36ac (Thermo Fisher, #39379), α-H3K36me3 (Abcam, ab9050). ChIP enrichment was quantified by qPCR with primers listed in Supplementary Table 5. For histone, ChIP values were normalized to H3. For H3K36me3 and H3K27me3, values were further normalized to enrichment at the positive control loci ACTIN for H3K36me3 and STM for H3K27me3.

### Micro Capture-C

Micro-Capture-C was performed as previously described by Hamley et al., 2023^10^ with modifications. 2.5 g of 10-day-old seedlings, as well as seedlings subjected to an additional 2 weeks vernalization treatment under constant 5 °C or daily FS conditions, were harvested and crosslinked for 10 min in 1% (w/v) formaldehyde in PBS. Crosslinking was quenched with glycine (0.125 M final concentration) for 5 min. Snap freeze with liquid nitrogen. Nuclei were extracted from 2 g of ground frozen material with Honda buffe and washed with Honda buffer three times. Subsequent steps were performed according to the protocol described by Hamley et al., 2023. Libraries were pooled and subjected to target enrichment using the myBaits Custom Kit under high-sensitivity hybridisation conditions. The sequence of all baits is listed in Supplementary Table 6).

### Single Molecule RNA FISH (smFISH)

The protocol described by Duncan et al., 2017^11^ was followed with minor modifications for root squash imaging. Briefly, root tips were fixed for 30 min in 4% paraformaldehyde (PFA), prepared from 16% EM-grade PFA (Electron Microscopy Sciences, 15710), and washed twice in 1× PBS. Roots were squashed onto slides under a coverslip and permeabilized in 70% ethanol (EtOH) for 4 h. Next, sequential 30 min incubations were carried out with 100% methanol (MeOH), then 100% EtOH followed by a 5 min incubation in wash buffer (2× SSC, Invitrogen, AM9763, containing 10% formamide (Sigma, F9037)). Slides were incubated overnight at 37 °C in the dark with Stellaris probes^12^ (Supplementary Table 7) conjugated with Quasar 670 dye (LGC Biosearch Technologies) diluted to a final concentration of 150 nM in hybridization solution (2× SSC, 10% dextran sulfate (Thermo Scientific, AC433240050), 10% formamide). The following day, samples were rinsed twice and incubated for 30 min in wash buffer. A second wash buffer containing 100 ng/mL DAPI (Thermo Scientific, 62248) was applied and samples were incubated for 30 mins at 37 °C. Slides were then washed in 2× SSC for 5 min and mounted in GLOX buffer (0.4% glucose, 10 mM Tris-HCl pH 8.0, 2× SSC) supplemented with glucose oxidase and catalase (Sigma-Aldrich; 1 µL each per 100 µL buffer).

Whole mount smFISH was performed as described by Zhao et al. 2023^13^ with minor modifications. Root tips were fixed as described above, then two brief 1 x PBS washes were carried out, followed by tissue permeabilization in 75% EtOH (∼16 h), 100% MeOH (2× 15 min), and 100% EtOH (2× 15 min), then a 1× PBS wash. The roots were cleared in ClearSee (10% (w/v) xylitol, 15% (w/v) sodium deoxycholate, 25% (w/v) urea)^14^ at 4 °C for 7 days then embedded in an acrylamide-based hydrogel (30% acrylamide-bisacrylamide (29:1), 10% ammonium persulfate, 5% TEMED). The roots were mounted on poly-L-lysine-coated slides with 20 µL embedding solution and allowed to polymerize for 20-30 min prior to coverslip removal. Hybridization and washes were performed as described above. Cell walls were stained with 0.1% (v/v) SCRI Renaissance 2200, followed by washing and mounting in 20 µL Vectashield (BioNordika, H-1000-10).

### Image Acquisition

Squashed-root imaging was performed using a Zeiss Elyra PS1 wide-field epifluorescence microscope equipped with a 100× oil-immersion objective (NA 1.46) and a cooled EMCCD camera (Andor iXon 897; 512 × 512 pixels, QE > 90%) operated in standard mode. Z-stacks spanning the entire nucleus were acquired with a step size of 0.2 µm. DAPI was excited at 405 nm and detected at 420-480 nm, while Quasar 670-labelled probes were excited at 642 nm and detected at 655-710 nm.

Whole-mount FISH imaging was performed using a Leica Stellaris 8 FALCON confocal microscope equipped with an HC PL APO CS2 63× water-immersion objective (NA 1.20) in photon-counting mode. Optical sections were acquired with a z-step of 0.3 µm at a scan speed of 400 Hz, with the pinhole set to 2 Airy units.

Quasar 670-labelled probes were excited at 645 nm using a white light laser (80 MHz, 85% laser power, 5% intensity), with emission collected at 660-730 nm (HyD X detector; line accumulation 6; frame averaging 2). SCRI Renaissance 2200 was excited at 405 nm, and emission was detected at 410-503 nm.

### Image Analysis

*COOLAIR* foci quantification was carried out using FIJI^15^. Volume and intensity were determined from root squash smFISH images with foci first saved as individual regions of Interest (ROIs). To determine volume, these ROIs were processed with a 3D Gaussian blur (σ = 0.5 µm in x, y, z) and background-subtracted (rolling ball radius = 20). Thresholding (2124-65535) was applied to generate masks, and volumes were quantified using the 3D Objects Counter plugin (minimum voxel count = 15, maximum = 19,488). To determine foci intensity, ROIs were blurred (σ = 0.5 µm), background-subtracted (rolling = 20), and summed along the z-axis. Thresholding was applied to select foci, and intensity values were measured using ROI Manager. Proportions of *COOLAIR* positive and negative cells were determined from z slices acquired around the mid-sections and epidermal regions of whole mount smFISH samples. Foci per nucleus data were determined from manually selected nucleus ROIs from maximum z-projections of these images using the Find Maxima command (prominence = 5), following smoothing (Gaussian blur, σ = 0.5 µm), and background-subtraction (rolling = 20).

### MNase-seq

MNase-seq was performed following the protocol described in Montez et al., 2025^16^. *ntl8-D3* seedlings were cultivated under warm conditions. Three biological replicates were used. For each replicate, 1.5 g of tissue was frozen in liquid nitrogen and ground into a powder. The material was resuspended in 35 mL of Nuclei Isolation Buffer (20 mM HEPES-KOH pH 7.4, 0.44 M sucrose, 1.25% Ficoll, 2.5% Dextran T40, 20 mM KCl, 0.5 mM spermidine, 5 mM DTT, 1% Triton X-100, protease inhibitors, and 0.1 mM PMSF). After filtration through two layers of Miracloth, the homogenate was centrifuged at 2000 × g for 15 min at 4 °C.

The nuclei pellet was washed once in 5 mL of isolation buffer and further purified using a Percoll gradient composed of 75% and 40% layers. Following centrifugation at 7000 × g for 30 min at 4 °C, nuclei were collected from the interface between the two layers and washed again in 6 mL of isolation buffer. Purified nuclei were resuspended in 1.5 mL MNase buffer (20 mM Tris pH 8.0, 0.125 M sucrose, 5 mM NaCl, 40 mM KCl, 2 mM CaCl_2_, protease inhibitors, and 0.1 mM PMSF), transferred to a clean tube, and pelleted at 1500 × g for 10 min at 4 °C. After removing the supernatant, the pellet mass was determined. A total of 65 mg nuclei in 300 μL MNase buffer was digested with 10 U MNase for 30 min at 37 °C. Reactions were terminated by adding EDTA, NaCl, SDS, and Proteinase K, followed by incubation at 65 °C for 1 h. DNA was purified by two phenol-chloroform extractions, one chloroform extraction, and ethanol precipitation with glycoblue at −80 °C. The DNA pellet was washed with 75% ethanol and resuspended in 30 μL water. RNA contamination was removed by RNase A treatment at 37 °C for 20 min. Mono- and subnucleosomal DNA fragments (< 200 bp) were enriched using AMPure XP beads while excluding fragments >300 bp. Sequencing libraries were prepared using the NEBNext Ultra II DNA Library Prep Kit according to the manufacturer’s protocol. After adaptor ligation, fragments > 200 bp were retained with AMPure XP beads, and libraries were amplified by 8 cycles of Illumina indexing PCR.

Libraries were pooled and subjected to target enrichment using the myBaits Custom Kit under high-sensitivity hybridisation conditions. The sequence of all baits is listed in Supplementary Table 6. Enriched libraries were sequenced on an DNBseq G400 platform (BGI Tech) with paired-end 150 bp reads. Data in Fig. S6 from FRI warm and cold has been re-plotted from published data in Montez et al., 2025^16^.

### Quant-seq

For Quant-seq experiments, total RNA was isolated as described for qPCR and further purified using the Qiagen RNeasy Miniprep Kit (74106). Sequencing libraries were prepared from 100 ng total RNA using the Quant-seq 3’ mRNA-Seq Library Prep Kit REV for Illumina (015UG009V0271) according to the manufacturer’s instructions. Library quality was assessed on a Fragment Analyzer using the HS-DNA assay, and concentrations were measured with a Qubit dsDNA HS assay (Thermo Fisher). Indexed libraries were pooled and sequenced on an Illumina NextSeq 2000 using a 100-cycle cartridge and the Lexogen Custom Sequencing Primer (CSP). Library preparation and sequencing were performed by Lexogen GmbH.

### Data processing and visualization

#### NET-seq bioinformatics analysis

The NET-seq data were obtained from previously published datasets^17^. To identify significant pausing sites, a sliding window approach was applied to scan signal enrichment across the COOLAIR gene region (step size = 50 bp; window size = 250 bp). For each window, the total signal within the window was defined as the observed value, while signals from the immediately upstream and downstream regions of equal window size were used as background controls. Statistical significance was assessed using a Poisson test and resulting *P*-values were adjusted for multiple testing using the Benjamini-Hochberg (BH) method.

#### tFLEP bioinformatics analysis

Processing of tFLEP-seq reads was performed using the pipeline available at https://github.com/ZhaiLab-SUSTech/FLEPSeq. Based on the presence or absence of poly(A) tails, reads were classified into non-poly(A) reads (representing elongating RNA) and poly(A) reads (representing polyadenylated RNA). To estimate Pol II occupancy, the number of reads corresponding to 3’ end positions was quantified and normalized against two housekeeping genes (*ACT7* and *UBC21*). For visualization, signal profiles were generated using a sliding window approach (step size = 20 bp; window size = 100 bp). To compare Pol II dynamics between conditions, three regions associated with major poly(A) clusters induced under cold treatment were defined. The proximal region was defined as ±300 bp surrounding the major poly(A) site (3,173,953) of the proximal isoform (Class I), while the distal region was defined similarly around the major poly(A) site (3,179,581) of the distal isoform (Class II). To avoid interference from proximal signals, a non-overlapping region (3,174,500-3,179,000) was defined as the gene body region. Detailed library quality metrics, including mapped rate, enrichment rate, proportion of poly(A), and proportion of non-poly(A) reads, are provided in Supplementary Table 8.

### Micro Capture-C sequencing bioinformatics analysis

Low-quality reads and adapter sequences were removed using fastp^18^. The resulting clean reads were aligned to the reference genome using BWA-MEM (https://github.com/lh3/bwa). PCR duplicates and reads with low mapping quality (MAPQ < 40) were filtered using pairtools^19^. Reproducibility and correlation between biological replicates were assessed using the *hicCorrelate* function in HiCExplorer^20^. Contact matrices were normalized by the total number of valid read pairs mapped within the selected *FLC* region and stored in *mcool* format using cooler^21^. The insulation score and boundaries within the FLC region were calculated at 1 kb resolution using cooltools ^22^. The observed/expected (O/E) matrix was calculated to correct for the distance-dependent decay of interaction frequency. Visualization of chromatin interaction patterns was performed using CoolBox^23^.

For the poly(A)-promoter contact analysis, the *FLC* locus was divided into 200-bp bins spanning the *FLC* gene region from 3,172,000 to 3,179,400, including the *COOLAIR* TSS region. Bins were classified as poly(A)-positive when the summed tFLEP-seq *COOLAIR* poly(A) read count across the three replicates exceeded 2 reads and as poly(A)-negative otherwise; promoter-contact OE values were then compared between the two classes using the AUC, calculated from the Mann-Whitney U statistic, with significance assessed by randomly reassigning the observed number of poly(A)-positive labels among bins and recalculating the AUC.

### Quant-seq analysis

Raw reads were quality-trimmed, and adapter sequences were removed using fastp^18^. Clean reads were aligned to the *Arabidopsis thaliana* reference genome (TAIR10) using STAR (https://github.com/alexdobin/STAR). The 3’ end sites were extracted from aligned reads by taking the terminal genomic position of each read in a strand-specific manner (the 3’ end of reads on the forward strand and the 5’ end of reads on the reverse strand). To examine antisense poly(A) pattern changes induced by FS, we analysed a set of genes with antisense transcription initiating close to their poly(A) sites, identified by plaNET-seq in a previous study^24^. For identification of poly(A) site clusters, sites on the same strand were sorted by genomic coordinate and a sliding-window algorithm was applied to merge adjacent sites within 24 bp into clusters. The highest-read-count site within the cluster was set as cluster summit. The clusters with a total read count below 5 were discarded.

## Acknowledgements

We thank Dr. Qianli Dong for help with Micro Capture-C and Shuqin Chen for excellent technical assistance, and current and former members of the Dean and Ding labs for helpful discussions. We thank Prof. Lori Passmore, Prof. Chang Liu, and Prof. Jiawei Wang for comments on the manuscript. We also thank the JIC Bioimaging facility and staff for their contribution to this publication.

## Author contributions

C.D. M.M. J.X, J.Z. & Y.D. conceived the study. J.X, C.G, M.S, E. M-B., M.Y, X.J, S.R., S.D., Y.Z., C.X., G-J.J. performed the experiments and analysed the data. G.M. analysed the data. C.D. obtained funding and C.D and Y.D supervised the work. J.X, C.G, G.M., M.M., C.D contributed to writing the manuscript and preparing the figures.

## Additional information

### Supplementary information

Supplementary files presenting individual replicate analysis of sequencing data and primer and probe information are provided with the manuscript.

